# Abnormal chondrocyte intercalation in a zebrafish model of *cblC* syndrome restored by an MMACHC cobalamin binding mutant

**DOI:** 10.1101/2023.01.20.524982

**Authors:** David Paz, Briana E. Pinales, Barbara S. Castellanos, Isaiah Perez, Claudia B. Gil, Lourdes Jimenez Madrigal, Nayeli G. Reyes-Nava, Victoria L. Castro, Jennifer L. Sloan, Anita M. Quintana

## Abstract

Variants in the *MMACHC* gene cause combined methylmalonic acidemia and homocystinuria *cblC* type, the most common inborn error of intracellular cobalamin (vitamin B12) metabolism. *cblC* is associated with neurodevelopmental, hematological, ocular, and biochemical abnormalities. In a subset of patients, mild craniofacial dysmorphia has also been described. Mouse models of *Mmachc* deletion are embryonic lethal but cause severe craniofacial phenotypes such as facial clefts. *MMACHC* encodes an enzyme required for cobalamin processing and variants in this gene result in the accumulation of two metabolites: methylmalonic acid (MMA) and homocysteine (HC). Interestingly, other inborn errors of cobalamin metabolism, such as *cblX* syndrome, are associated with mild facial phenotypes. However, the presence and severity of MMA and HC accumulation in *cblX* syndrome is not consistent with the presence or absence of facial phenotypes. Thus, the mechanisms by which mutation of *MMACHC* cause craniofacial defects have not been completely elucidated. Here we have characterized the craniofacial phenotypes in a zebrafish model of *cblC* (*hg13*) and performed restoration experiments with either wildtype or a cobalamin binding deficient MMACHC protein. Homozygous mutants did not display gross morphological defects in facial development, but did have abnormal chondrocyte intercalation, which was fully penetrant. Abnormal chondrocyte intercalation was not associated with defects in the expression/localization of neural crest specific markers, *sox10* or *barx1*. Most importantly, chondrocyte organization was fully restored by wildtype MMACHC and a cobalamin binding deficient variant of MMACHC protein. Collectively, these data suggest that mutation of *MMACHC* causes mild to moderate craniofacial phenotypes that are independent of cobalamin binding.

## Introduction

Methylmalonic aciduria and homocystinuria, cblC type (*cblC*) (MIM 277400) is caused by mutations in the *MMACHC* gene (Lerner-Ellis et al., 2006). *cblC* is characterized by heterogenous phenotypes spanning cobalamin deficiency, microcephaly, failure to thrive, hydrocephalus, neurological abnormalities, hematological manifestations, ocular deficits, and craniofacial dysmorphia (Carrillo-Carrasco et al., 2012a; Martinelli et al., 2011a). Craniofacial phenotypes associated with *cblC* include a long face, high forehead, flat philtrum, and large low-set ears, which have been reported in approximately 20% of patients (Biancheri et al., 2001; Carrillo-Carrasco et al., 2012b; Cerone et al., 1999; D’Alessandro et al., 2010a; Fischer et al., 2014; Martinelli et al., 2011b; S et al., 2014). There are multiple subtypes of cobalamin deficiency and additional subtypes have also been associated with craniofacial phenotypes (Bassim et al., 2009; Coelho et al., 2012, p. 4; Gérard et al., 2015, p. 1; Oladipo et al., 2011). For example, *cblX* syndrome (MIM 309541), an X-linked subtype, is associated with craniofacial abnormalities (Yu et al., 2013). *cblX* does not result from mutations in the *MMACHC* gene but is associated with reduced *MMACHC* expression which disrupts cobalamin metabolism. The presence of facial phenotypes across multiple cobalamin subtypes suggests that defects in cobalamin processing/metabolism are a potential cause for abnormal facial development.

*MMACHC* encodes an enzyme with decyanase and dealkylase activity (Froese et al., 2012). *cblC* mutations are thought to disrupt cobalamin binding and/or enzyme activity reducing the synthesis of adenosylcobalamin and methylcobalamin, which are cofactors for the enzymes methylmalonyl-CoA mutase and methionine synthase, respectively (Carrillo-Carrasco et al., 2012b). This leads to the accumulation of methylmalonic acid and homocysteine. Interestingly, *cblX* syndrome results in atypical craniofacial development in the presence of mild or in some cases completely absent cobalamin phenotypes (Huang et al., 2012; Koufaris et al., 2016). Therefore, the possibility exists that *MMACHC* may regulate craniofacial development outside of its characterized cobalamin binding and enzymatic roles.

Morpholino mediated knockdown of *mmachc* in zebrafish was the first to describe dramatic defects in facial development (Quintana et al., 2014). These included loss of ceratobranchial cartilages and severe truncation of the Meckel’s cartilage with inversion of the ceratohyal. These data are consistent with a zebrafish model of *cblX* syndrome (morpholino mediated knockdown), which demonstrates overlapping craniofacial phenotypes. Interestingly, facial phenotypes in the zebrafish model of *cblX* syndrome were not associated with detectable levels of cobalamin deficiency but were restored by ectopic expression of *MMACHC* (Quintana et al., 2014). On the surface, these data suggest that in *cblC* and *cblX*, abnormal expression/function of MMACHC is the cause of abnormal craniofacial development. However, facial phenotypes in a knock-in mouse model of *cblX* syndrome cannot be restored by cell type specific expression of *Mmachc* (Chern et al., 2022). These data are particularly confounding because a mouse model of *cblC* displays facial clefting and fusion of the palate, which was reversed by overexpression of MMACHC (Chern et al., 2020). Based on these studies, the function of MMACHC in facial development are still unresolved.

Deletion of *Mmachc* in mice is embryonic lethal, which has complicated the early analysis of facial development (Chern et al., 2020). Given the lethality of mouse models, a zebrafish germline mutant (hg13 allele) was recently created (Sloan et al., 2020). Although homozygous mutation in zebrafish was lethal during the juvenile period, the hg13 allele displayed the characteristic growth impairment, a retinal phenotype, and elevated methylmalonic acid (MMA) by 7 days post fertilization (DPF) (Sloan et al., 2020). Other metabolites such as homocysteine were not measured and undetectable. Our analysis of the hg13 allele demonstrated mild craniofacial abnormalities characterized by an increased extension of Meckel’s cartilage from the tip of the ceratohyal, increased width of the palatoquadrates, and abnormal chondrocyte intercalation in the hyosymplectic cartilage. These mild phenotypic changes did not alter the angle between specific cartilage structures of the viscerocranium, allowing development of an overall normal facial skeleton. We did not detect changes in expression or location of neural crest cell (NCC) markers, *sox10* or *barx1*, at 1 and 2 DPF. Defects in chondrocyte development were restored via global expression of an *MMACHC* transgene. These data indicate that *mmachc* directly regulates chondrocyte development. Most fascinating was the fact that site directed mutagenesis of the MMACHC cobalamin binding domain did not preclude restoration of the phenotypes present in the hg13 allele. Collectively, these data suggest that *mmachc* regulates craniofacial development independent of cobalamin binding.

## Materials and Methods

### Zebrafish

For all experiments, embryos were obtained by natural spawning using adult *mmachc*^*hg13/+*^, *Tg*(*sox10:tagRFP*) carrying the hg13 allele, *Tg*(*ubi:MMACHC*) with or without the hg13 allele, or *Tg*(*col2a1a:EGFP*) carrying the hg13 allele. All embryos were maintained in E3 media at 28°C with a 14/10 light: dark cycle. All adults and larvae were maintained according to guidelines from The University of Texas El Paso Institutional Animal Care and Use Committee (IACUC). Euthanasia was performed according to guidelines from the 2020 American Veterinary Medical Association and based on approved protocol 811869-5. The *Tg*(*ubi:MMACHC*) allele was produced using gateway cloning. The *MMACHC* open reading frame, previously described (Quintana et al., 2014) was cloned into pME-MCS using BamHI and SacII cloning sites. An LR cloning reaction was then utilized to produce the *Tg*(*ubi:MMACHC*) in the pDestTol2CG2 backbone with p5e-Ubiquitin and p3ePolyA. The destination vector was injected at the single cell stage as described (Kwan et al., 2007) and genotyping was performed by detecting human *MMACHC* using Fwd: ACTTACCGGGATGCTGTGAC and Rev: GGGTGTGGTAAAGGGAAGGT primers.

### Genotyping

Genotyping was performed using a nested PCR approach combined with restriction enzyme digest. DNA was obtained by excising a small piece of the developing tail and collected in individual PCR tubes. Samples were heated for 5min at 95°C and cooled to 4°C for 10min. and then 6μL of 500mMTris, pH:8.0 (Fisher Scientific) solution was added to each tube. Samples were centrifuged for 10 minutes at 10,000Xg. Lysate was used directly for PCR (2μl). The following primers were designed to amplify the region flanking the *mmachc* hg13 allele:exon2: flankingPCR1 5’-TACTGGTCTCCGAGAGGGACTG-3’, flankingPCR1 5’-CAGAGAGATGCAGGCAGACA-3’, FWPCR2 5’-TACTGGTCTCCGAGGGACTG-3’, RVPCR2 5’-TGTAAGAAGGGCAGGAATGC-3’. PstI (New England Biolabs) restriction enzyme digest was performed after amplification to identify carriers. Digestion was performed according to manufacturer’s protocol. The hg13 allele abrogates PstI digestion and therefore, can be used to detect the genotype of interest.

### In situ hybridization

*barx1* RNA probe was cloned into the pGEM T-easy vector and transformed into *E. coli* competent cells. Amplification of the *barx1* coding sequence was performed with the following primers: *barx1* FWD 5’-GAGATTGGGGCGCACTATTA-3’ and *barx1* REV 5’-GCACGGATCGGGATAATCTA-3’. Target region was cloned into the pGEM T-Easy vector and sequence validated. PCR was used to amplify target region from the pGEM vector with FWD: 5’-GTTTTCCCAGTCACGACGTT-3’ and T REV: 5’-TTTATGCTTCCGGCTCGTAT-3’ primers. RNA probe synthesis was performed using purified PCR product as described (Quintana et al., 2014). *In situ* hybridization was performed as described with minor modification (Thisse and Thisse, 2008). Zebrafish embryos were dechorionated and collected at 48 hours post fertilization (HPF). Embryos were fixed in 4% paraformaldehyde (Electron Microscopy Sciences) overnight at 4°C. The next day, embryos were rinsed in 1x phosphate buffered saline (PBS) (Fisher Scientific) before removal of pigmentation. Pigment was removed by bleaching embryos with bleach solution (3% hydrogen peroxide/0.5% potassium hydroxide) (Fisher Scientific). Following depigmentation, embryos were dehydrated in the following series: 25%(vol/vol), 50%(vol/vol), and 75%(vol/vol) methanol: phosphate buffered saline (PBS) (Fisher Scientific) and stored at -20°C overnight in 100% methanol. Prior to hybridization protocol, embryos were rehydrated in successive dilutions of 75%, 50%, and 25% methanol: PBS (Fisher Scientific). Prior to permeabilization, embryos were washed four times in 100% 1x PBS with 0.1% Tween 20 (PBT). Permeabilization was performed with proteinase K (10μg ml^-1^ in PBT) for 30 minutes at room temperature and stopped with 4% paraformaldehyde for 20 minutes. Permeabilized embryos were prehybridized by adding 500μL of hybridization mix (50% deionized formamide, 5x saline sodium citrate (SSC), 0.1% Tween 20, 50μg ml^-1^ of heparin, 500μg ml^-1^ of RNase-free tRNA adjusted to pH 6.0 by adding citric acid for 2 hours at 70°C. After 2 hours of incubation, 1.5 μL of antisense (50μg/μL) and 1.5μL of sense (50μg/μL) probes were incubated overnight at 70°C. Post hybridization, samples were gradually rinsed in 75%(vol/vol), 50%, 25% and 100% hybridization buffer: 2x SSC (Fisher Scientific) buffer for 10 minutes at 70°C. Two final washes for 30 minutes in 0.2x SSC at 70°C were performed. Gradual conversion of each sample to 1XPBT was performed using a diluted series of SSC/PBT and samples were blocked in blocking buffer for 3 hours at room temperature before adding 1:10000 of anti-DIG antibody (Millipore/Sigma) overnight at 4°C. All samples were developed with BM purple 4°C and imaged on a Zeiss Stereo Microscope at 8X magnification.

### In vitro mRNA synthesis and microinjection

*MMACHC* (RefSeq NM_015506.2) mRNA was synthesized with the T7 Ultra mMessage mMachine kit (Fisher Scientific) according to manufacturer’s instructions and from a commercially available vector previously described (Quintana et al., 2014). Missense mutation was introduced into the open reading frame using the QuikChange Site Directed mutagenesis kit (c.440G>A) (Fisher Scientific) according to manufacturer’s instructions. Primers were as follows: Fwd: GAGAGAGCCTCCCAGAGCTACAGATAGAAATCATTGCTG and Rev: CAGCAATGATTTCTATCTGTAGCTCTGGGAGGCTCTCTC. Synthesized RNA was injected at a final concentration of 800ng RNA/embryo at the single cell stage according to previously described empirical derivation (Quintana et al., 2014). Larvae were maintained in embryo media at 28.5°C until 5 DPF and fixed in 4% paraformaldehyde (Fisher Scientific) at room temperature for 1 hour and stored in 1x PBS (Fisher Scientific) before analysis.

### Alcian blue staining and microscopy

Alcian blue was performed at 5 DPF as previously described (Quintana et al., 2014). High resolution images were taken after dissection of the viscero- and neurocranium elements using a Leica Laser Micro-dissection microscope at 40 and 63X magnification. Representative images were taken using a 1 × 1 grid. For confocal microscopy, transgenic *hg13* larvae (*mmachc*^*hg13/hg13*^ *Tg(col2a1a:EGFP*)) were staged and fixed at 5 DPF with 4% paraformaldehyde (Fisher Scientific). Larvae were dissected for genotyping and unbiased imaging was performed on the remaining head tissue. For mounting of the hyosymplectic, the eyes were excised from craniofacial region. Mutant and wildtype heads were angled along either a left or right ventrolateral angle and mounted in glass bottom dishes (Fisher Scientific) using 0.6% low-melt agarose (Fisher Scientific). Confocal imaging was performed on a Zeiss LSM 700 at 40X oil magnification. Images were restricted to the larval craniofacial region, particularly around the hyosymplectic region. For each fish, a minimum of 8 to 20 z-stacks were collected. For confocal microscopy of NCC markers, transgenic *hg13* larvae (*mmachc*^*hg13/hg13*^ *Tg (sox10:RFP))* were staged and fixed at 30 and 40 HPF. Larvae were mounted along a right lateral angle in glass bottom dishes (Fisher Scientific) using 0.6% low-melt agarose (Fisher Scientific). Confocal images were performed on a Zeiss LSM 700 microscope at both 20X and 40X oil magnification. For each fish at 20X, a minimum of 34 to 57 z-stacks were collected. For 40X, a minimum of 33 to 55 Z-stacks were collected. Maximum intensity projection images were used to analyze the formation of the pharyngeal arches as shown.

### Image quantification, chondrocyte patterning, and whole head angle measurements

Fiji, an open-source platform (Schindelin et al., 2012) was utilized to measure the angles between individual chondrocytes within the hyosymplectic region. A Fire LUT was applied using ImageJ software to provide a fluorescent intensity gradient with equal pixel (0 to 255) measurements to ensure the angles were present in the same z-plane to avoid bias. The posterior portion of the hyosymplectic area was evaluated for abnormal chondrocyte intercalation. The angle tool was implemented to draw at the center of 3 adjacent cells along the region of interest. 5 different measurements for 3 individual/adjacent chondrocytes were recorded for each subject. Methods for angle measurements were performed as previously described (Shull et al., 2022). Total chondrocyte measurements from *hg13-/-* (n=41) and sibling (n=30) were averaged and plotted on a bar graph for comparison. Additional whole morphology head angle measurements were performed (Staal et al., 2018). Angles between structures included the Meckel’s cartilage and palatoquadrate (M-PQ), the angle of the ceratohyal (CH), angle between the palatoquadrate and ceratohyal (PQ-CH), and the angles Meckel’s (M) angle. Examples of each measurement are shown in Figure 1. In addition, the length of the Meckel’s cartilage from the tip of the ceratohyal and the width between the palatoquadrates was measured as shown in Figure 1C (Quintana et al., 2017a). Statistical significance was performed using a standard T-test.

**Figure 1.**
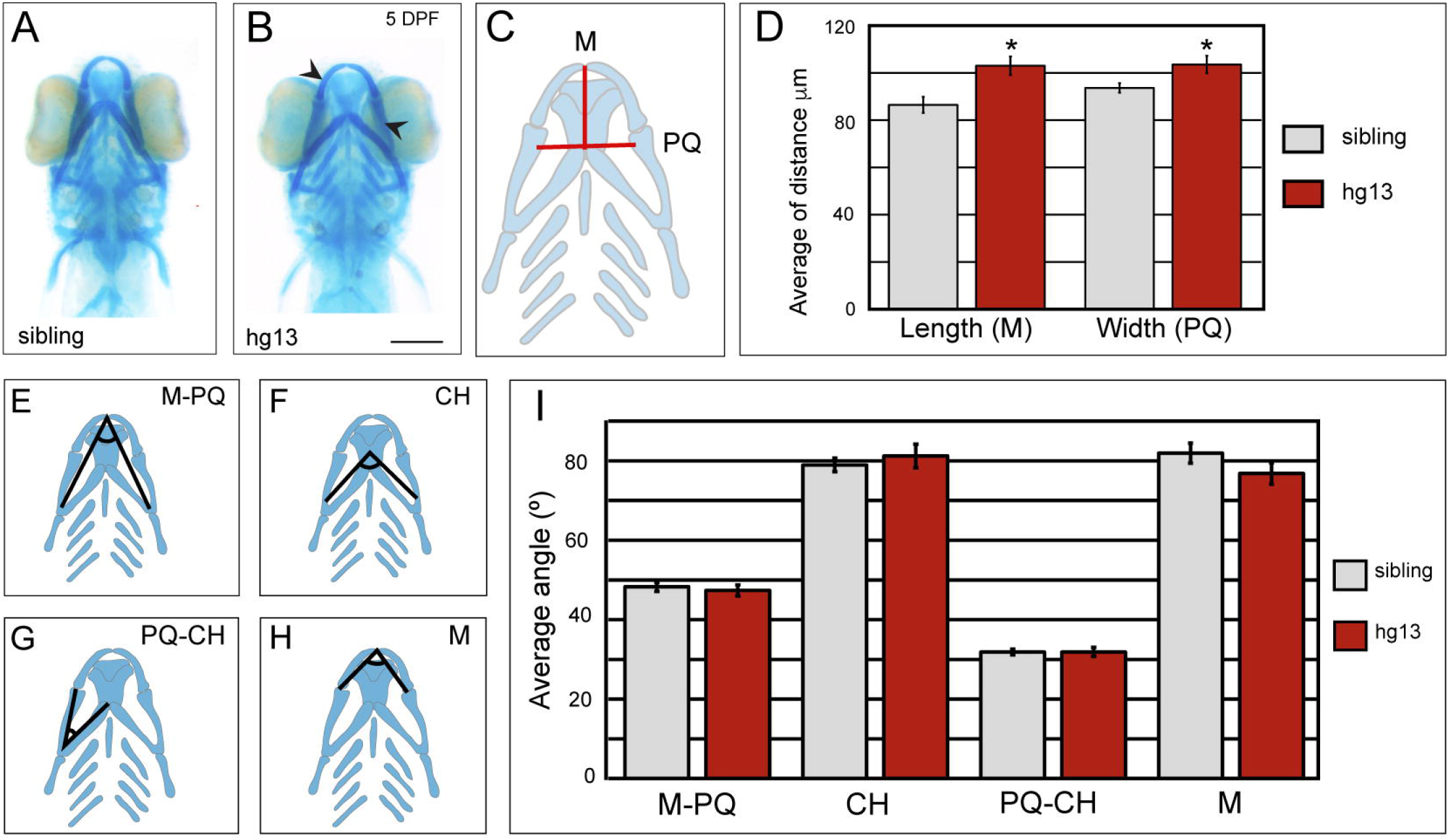
Mutation of *mmachc* affects extension and width of the Meckel’s cartilage. A-B. Wildtype siblings (sibling) and hg13^-/-^ mutants (hg13) were stained with alcian blue at 5 days post fertilization (DPF) to analyze the developing viscerocranium. Arrows indicate regions of increased width (PQ) and length (M). C. Schematic to illustrate location of length (M) and width (PQ) measurements. D. Quantification of the length from the Meckel’s cartilage (M) to the top of the ceratohyal (CH) and the width from the inner most region of either side of the palatoquadrates (PQ). N= 19, hg13; N= 14, sibling. p<0.05. Scale bar 200 μm; magnification at 8x. E-H. Schematic diagrams indicating angle measurements collected between structures of the viscerocranium. Measurements include the Meckel’s cartilage to the palatoquadrate (M-PQ), angle of the ceratohyal (CH), palatoquadrate to the ceratohyal (PQ-CH), and the Meckel’s cartilage angle (M). I. Quantification of angle measurements performed at each of the regions indicated in E-H. No significant difference was detected in any of the regions analyzed. N= 41, hg13; N= 30, sibling

## Results

### *mmachc* mutants do not show overt morphological defects in craniofacial development

Previous studies suggest that global deletion of *Mmachc* in mice causes craniofacial abnormalities (Chern et al., 2022, 2020). We therefore sought to characterize craniofacial development in a zebrafish germline mutant (hg13 allele) of *mmachc*. The hg13 allele was previously validated and ocular phenotypes were characterized, but craniofacial phenotypes have yet to be studied in this model (Sloan et al., 2020). We stained larvae at 5 DPF with alcian blue to study cartilage development from homozygous mutants of the hg13 allele and their wildtype clutch mates. Initial qualitative observations did not demonstrate overt morphological defects in homozygous animals (Figure 1A&B). However, since craniofacial phenotypes are mild to moderate in *cblC*, we sought additional quantitative measurements of facial development (Quintana et al., 2017a). We measured the length from the ceratohyal to the tip of the Meckel’s cartilage (M) and the width between the palatoquadrates (PQ) (Figure 1C: red lines). We observed an increased width between each end of the palatoquadrates as described in Figure 1D and an increased extension of the Meckel’s cartilage from the ceratohyal (Figure 1D, p<0.05). We then employed angle measurements between specific structures of the viscerocranium as previously described (Staal et al., 2018). We measured the angle between the Meckel’s cartilage: palatoquadrate: M-PQ (Figure 1E), the angle of the ceratohyal: CH (Figure 1F), the palatoquadrate: ceratohyal angle: PQ-CH (Figure 1G), and angle of the Meckel’s cartilage: M (Figure 1H) in sibling wildtype and the hg13 allele. We did not detect any significant differences in the formation of these structures in mutant larvae (Figure 1I). Despite the normality of the angles between these cartilage structures, we noted that increased extension of the Meckel’s cartilage has been used as an indicator of facial clefting in previous studies (Quintana et al., 2017a).

### Mutation of *mmachc* is associated with abnormal chondrocyte intercalation

To determine if abnormal cartilage development caused subtle changes in the extension of the Meckel’s cartilage and width of the palatoquadrate, we performed alcian blue staining at 5 DPF and performed high resolution imaging of the palatoquadrate and adjacent hyosymplectic structure (Figure 2A&B). While chondrocytes within the palatoquadrate were stacked adjacent (penny like stacking) to one another in wildtype siblings (Figure 2A-A’), chondrocytes from homozygous mutant larvae were disorganized and displayed an abnormal pattern as shown in Figure 2B-B’. We further characterized these deficits in chondrocyte development using the *Tg*(*col2a1a:EGFP*) transgene. High resolution confocal microscopy demonstrated normal chondrocyte organization in wildtype clutch mates at 4 and 5 DPF (Figure 2D&G). In contrast, we noted abnormal chondrocyte intercalation in the ventral end of the hyosymplectic at 4 DPF in homozygous mutants (Figure 2E-E’). The phenotype was exacerbated at 5 DPF whereby both the ventral and dorsal (Figure 2F, see shaded areas for dorsal ventral nomenclature) hyosymplectic regions were abnormally organized (Figure 2H-H’’). We measured the angle between adjacent nuclei (3 adjacent cells with 5 measurements/larvae) as shown in Figure 2I schematic. The average angle between nuclei in sibling wildtypes was 156.9°, but the average angle between chondrocytes of homozygous mutants was 121.2° (p<0.05) (Figure 2I: bar graph). These data provide evidence that *mmachc* regulates chondrocyte organization/intercalation.

**Figure 2.**
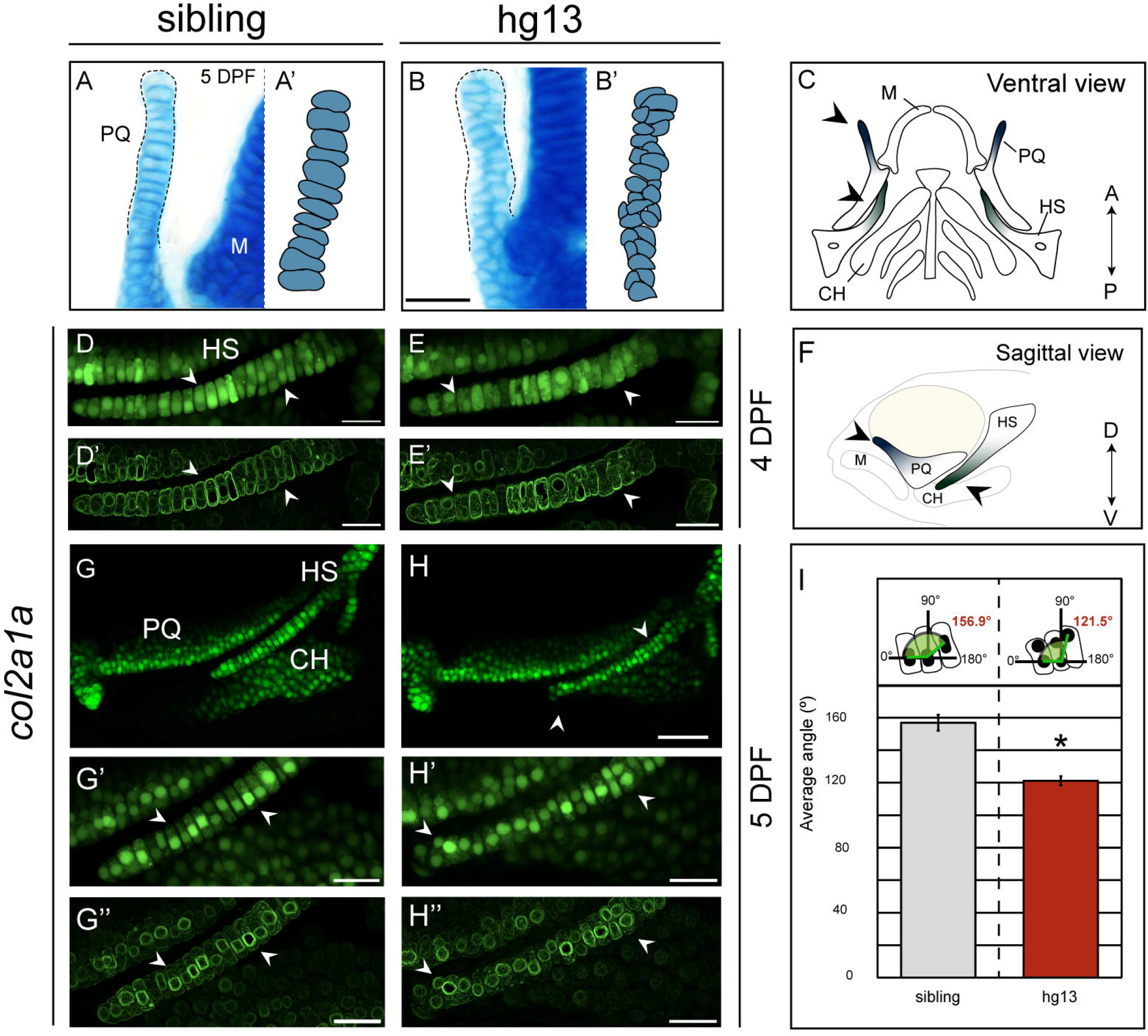
Abnormal chondrocyte organization observed in the hg13 allele. A-B. Alcian blue was performed at 5 days post fertilization (DPF) on sibling and hg13^-/-^ (hg13) mutant larvae. Based on distance measurements in Figure 1, high resolution imaging of the palatoquadrate (PQ) was performed. Dotted lines demarcate the region where differences in skeletal elements were observed. Digital drawings in panel A’ and B’ are provided for clarity. Magnification at 63X; scale bar at 30 μm. Numbers of total animals hg13, N= 19; sibling, N= 14. C. Ventral schematic of the viscerocranium with colored gradients in the PQ and hyosymplectic (HS) where aberrant organization was observed. Meckel’s cartilage (M) and ceratohyal (CH) are indicated for clarity. The dorsal extension of the PQ (blue gradient) represents the area observed by alcian blue stain (A&B) and the ventral extension of the HS is indicated (green gradient) D–H. Confocal microscopy of sibling and hg13 larvae harboring the *Tg*(*col2a1a:EGFP*) reporter at 4 & 5 DPF. Uniform stacking is shown in panel D&G of sibling (4 & 5 DPF, respectively). In contrast, abnormal stacking is observed in the HS of hg13 mutants (E&H); arrows indicate areas of abnormal intercalation. Magnification at 40X; scale bar at 20 μm (D&E; G’&H’) and 50 μm (D’&E’; G&H). Panel F provides a sagittal schematic for clarity, with HS and PQ highlighted. D’&E’;G’’&H’’. Images from D&E;G&H were manipulated using the edges plugin from ImageJ software. Irregularities are observed in chondrocyte intercalation and nuclear angle. I. The angle between 3 adjacent nuclei was quantified. Five independent measurements per larvae were performed. The average angle between 3 adjacent cells was quantified and plotted. hg13 N= 30; sibling N=30, p<0.05. Abbreviations: (A) anterior, (P) posterior, (D) dorsal, (V) ventral.

### Early neural crest cell (NCC) development is unchanged by mutation of *mmachc*

Cranial NCCs produce many of the cartilage structures of the viscerocranium. We analyzed *sox10* expression and location in the pharyngeal arches at 30 HPF using the *Tg*(*sox10:TagRFP*) reporter. Sox10 is a marker of NCCs (Honoré et al., 2003) and pharyngeal arches are transient embryonic structures that produce cartilage and bone. We observed 5 complete pharyngeal arch structures in both sibling and hg13 homozygous mutants (Figure 3A-B & A’-B’) indicating proper formation of the pharyngeal arch structures. We noted cartilage defects at 4 and 5 DPF (Figure 2) and therefore analyzed additional timepoints of NCC development. We measured expression of *sox10* and *barx1* at 40 HPF and 2 DPF, respectively. Barx1 is required for cranial NCC development (Sperber and Dawid, 2007). We did not detect severe disruptions in the expression or location of *sox10* at 40 HPF (Figure 3C-D and 3C’-D’), which was consistent with normal expression and location of *barx1* expression at 2 DPF (Figure 3E-F). *barx1* expression was analyzed by in situ hybridization and the area of *barx1* expression was quantified using pixel intensity (Figure 3G). No significant difference was observed.

**Figure 3.**
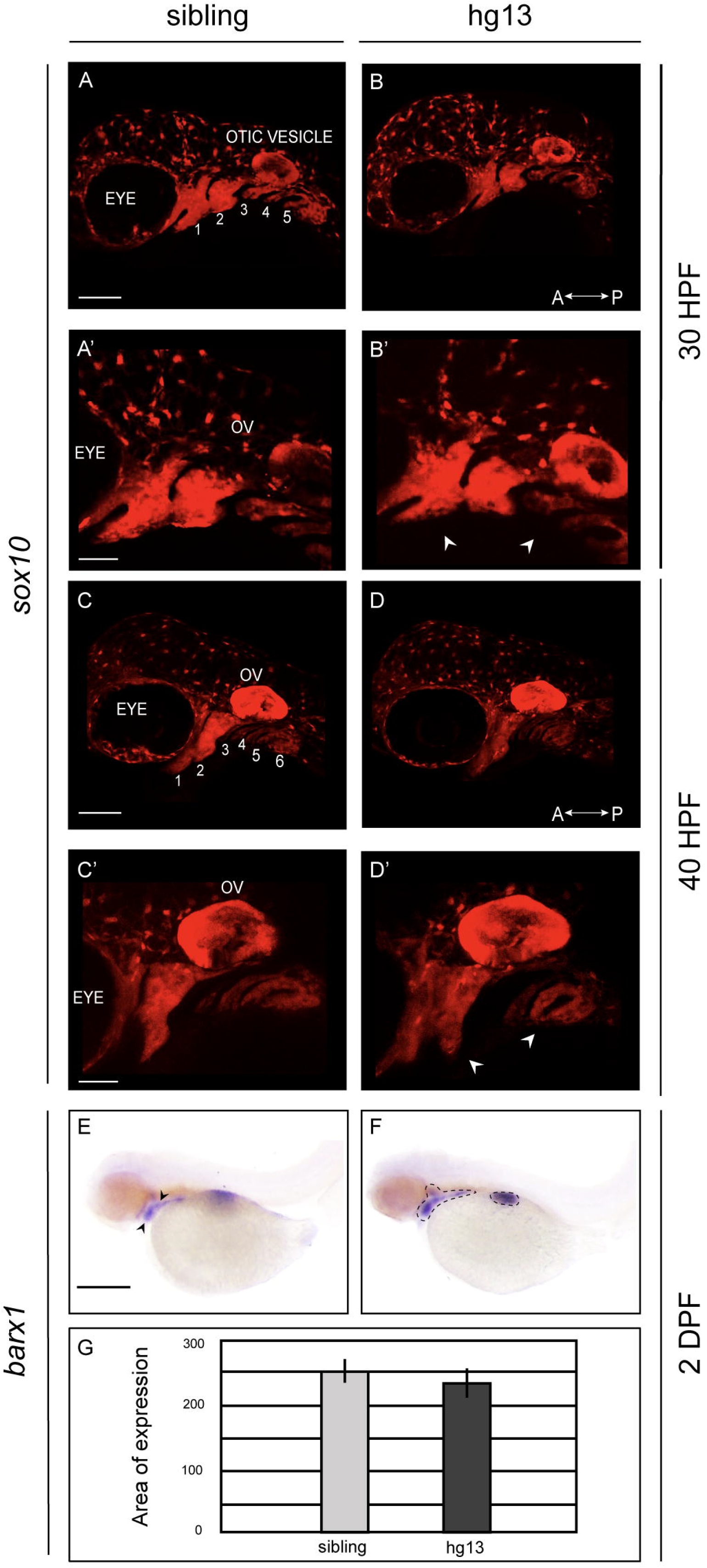
Early neural crest cell markers revealed no changes in expression. A-D. Confocal microscopy (20X magnification: Scale bar at 100 μm) was performed to view the developing pharyngeal arches in siblings and hg13^-/-^ mutants (hg13) carrying the *Tg*(*sox10:TagRFP*) reporter at 30 (A&B) (30 HPF hg13 N=6; sibling N=5) and 40 (C&D) (40 HPF hg13, N=2; sibling, N=4) hours post fertilization (HPF). A’−D’. 40X magnification of A−D with scale bar of 50 μm. Regions labeled for clarity are the eye and otic vesicle with each arch numbered. Arrows indicate pharyngeal arch structures. Anterior (a) and posterior (p) are indicated. E-F. Whole mount *in situ* hybridization (ISH) with *barx1* specific probed performed at 2 days post fertilization (DPF). Scale bar 200 μm magnification at 8X. Dotted lines demarcate areas with *barx1* expression at 2 DPF in sibling (E) and (F) hg13 mutant. G. Quantification of pixel intensity in E&F using ImageJ software. hg13 N=4; sibling N=3. Abbreviations: (OV) otic vesicle, (A) anterior, (P) posterior

### Cobalamin binding is not required for normal chondrocyte stacking

Facial phenotypes are not completely penetrant in *cblC*, but cobalamin deficits are. Therefore, we asked whether cobalamin binding/processing is essential for proper facial development. We designed a restoration experiment whereby we injected mRNA encoding *MMACHC* harboring the patient derived mutation p.Gly147Asp. The p.Gly147Asp variant is in the cobalamin binding domain of MMACHC. Previous studies have demonstrated that this variant completely abolishes the ability of MMACHC to bind to both cyanocobalamin and hydroxycobalamin, eliminating enzymatic activity, and resulting in early onset cobalamin deficits (Froese et al., 2009). We injected RNA encoding the MMACHC^p.Gly147Asp^ variant into hg13 mutants and their clutch mates to subsequently analyzed chondrocyte intercalation/organization at 5 DPF. We again observed abnormal chondrocyte intercalation in the non-injected hg13 homozygous larvae within the hyosymplectic region (Figure 4A-B and 4A’-B’). This resulted in a decreased angle between nuclei consistent with data in Figure 2 (Figure 4D-E, gray and dark gray bars). As a validation measure we used Fiji, an open-source platform (Schindelin et al., 2012), to apply a pseudocolor “Fire” lookup table (LUT), to distinguish differences in light intensity. This approach circumvents any bias when measuring through the z-plane. The pseudocolor was used as a control to perform additional angle measurements and ensure that all cells measured were indeed within the same Z-plane (Figure 4A”-B”). Our analysis demonstrated reduced overall angles between cells as was detected using manual quantification (Figure 2) as described in the material and methods section. Intriguingly, the injection of MMACHC^p.Gly147Asp^ encoding mRNA into hg13 homozygous embryos was able to restore the angles between nuclei to normal levels as shown in Figure 4C-C” and quantified in Figure 4D (p<0.05). Importantly, we performed restoration of the hg13 chondrocyte phenotype using the *Tg*(*ubi:MMACHC*) allele (ute2tg), which expresses wildtype MMACHC under the control of the ubiquitin promoter. This assay served as a control experiment, with which to test the specificity of the restoration and phenotype. As shown in Figure 4E, the ubiquitous expression of wildtype *MMACHC* completely restored the angles between nuclei and chondrocyte intercalation (p<0.05).

**Figure 4.**
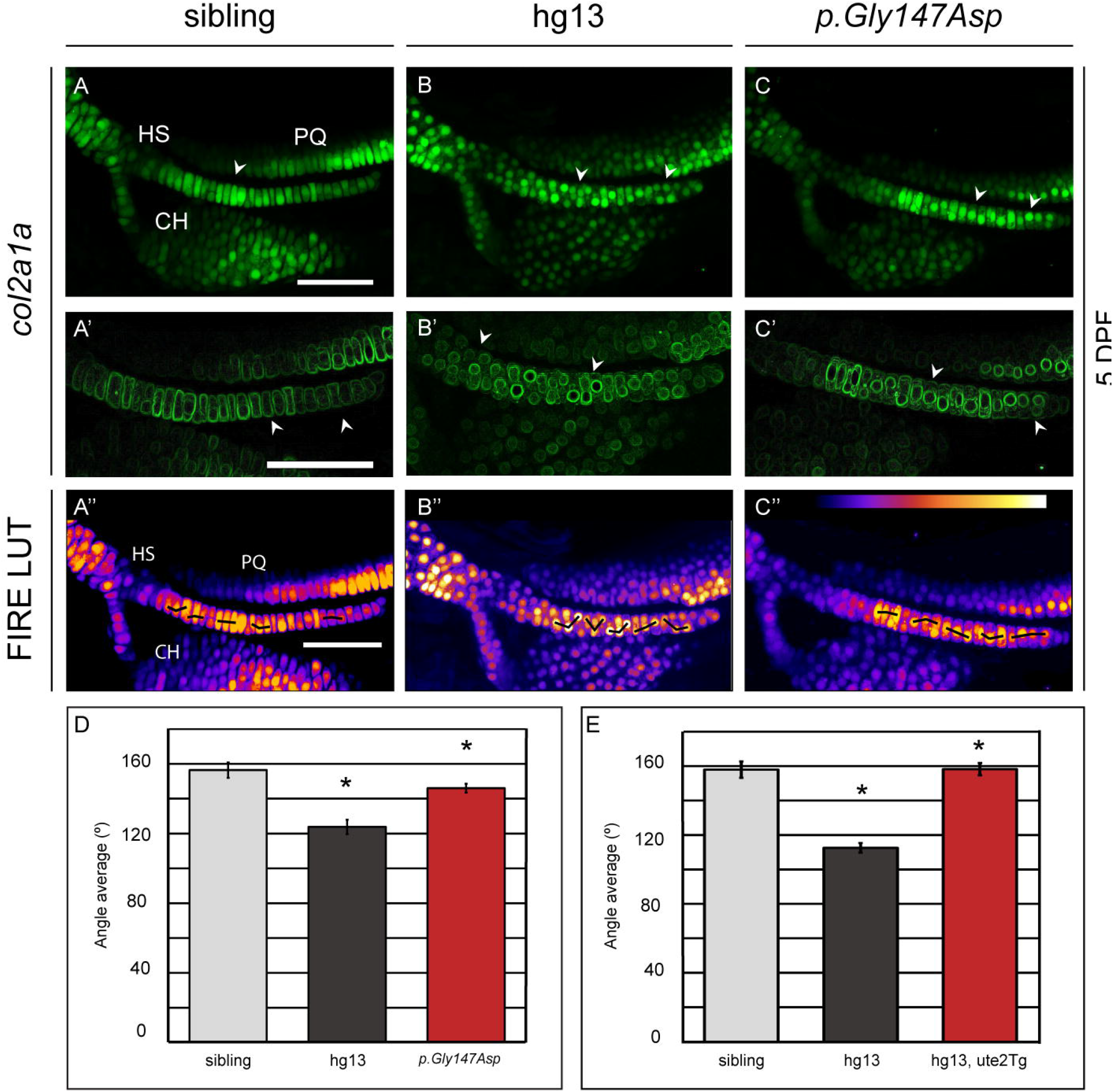
Chondrocyte stacking occurs independent of cobalamin binding. A–C. Sibling or hg13 homozygous mutants (hg13) carrying the *Tg*(*col2a1a:EGFP*) were injected at the single cell stage with mRNA encoding the p.Gly147Asp *MMACHC* variant. A-B are non-injected clutch mates to those injected (p.Gly147Asp). Confocal microscopy was used to analyze chondrocyte intercalation in the hyosymplectic (HS) and palatoquadrate (PQ) at 5 days post fertilization (DPF). sibling, N=7; non-injected hg13, N=7; MMACHC^p.Gly147Asp^, hg13 N=7; Magnification at 40X; scale bar at 50 μm. p<0.05. A’–C’. Images from A-C were modified using the edges plugin from ImageJ software to visualize nuclei. A”–C”. A pseudocolor LUT was applied to images from A–C to provide a color map of high and low light intensities (highest to low signal: yellow, red, purple; 0-255) throughout a z-stack image. D-E. Quantification of angle measurements as shown by the black lines in A”C”. For experiment described in (E), sibling (N=5), hg13 (N=5), hg13 ute2Tg (carrying the *Tg*(*ubiquitin:MMACHC*) transgene) (n=6), p<0.05. Abbreviations: (CH) ceratohyal

## Discussion

Mutation of *MMACHC* in humans causes *cblC* disorder. Mild to moderate facial dysmorphia has been observed in a subset of patients (Biancheri et al., 2001; Carrillo-Carrasco et al., 2012a; Cerone et al., 1999; D’Alessandro et al., 2010b; Martinelli et al., 2011a) but these phenotypes are not well characterized in humans, are not completely penetrant, and can be subtle in nature. Interestingly, other subtypes of cobalamin deficiency have also been associated with facial phenotypes (Bassim et al., 2009; Coelho et al., 2012, p. 4; Gérard et al., 2015, p. 1; Oladipo et al., 2011). For example, a zebrafish model of *cblX* syndrome is associated with craniofacial deficits, which can be restored by exogenous expression of MMACHC (Quintana et al., 2014). However, the mechanisms by which MMACHC expression regulates facial development are not completely characterized. Here we characterized the cellular phenotypes associated with mutation of zebrafish *mmachc*. We questioned whether MMACHC cobalamin binding activity is required for proper facial development. Interestingly, we restored facial phenotypes in the hg13 allele through expression of a cobalamin binding mutant know to cause *cblC*. Collectively, these data implicate MMACHC in the regulation of craniofacial development and suggest that a domain outside of the MMACHC cobalamin binding domain is required for facial development.

We observed nearly normal morphology of cartilage structures in the hg13 allele, but we identified a highly penetrant defect in chondrocyte intercalation in the palatoquadrate and hyosymplectic region. Interestingly, *barx1* and *sox10* expression was normal during early NCC development suggesting that the neural crest undergoes proper specification and migration. Similar phenotypes have been reported in glypican 4 mutants (Sisson et al., 2015) whereby defects in chondrocyte intercalation are present but neural crest markers are normal. We specifically observed abnormal intercalation in the palatoquadrate and hyosymplectic, which are derived from pharyngeal arch 1 and 2, respectively (Birkholz et al., 2009). Pharyngeal arch 2 cartilage elements begin intercalation at approximately 50–54 HPF (Clément et al., 2008) and consequently our analysis at 5 DPF, demonstrates the near final organization of these cells. Notably, we observed abnormal intercalation at 4 DPF, but only in the ventral most region of the hyosymplectic, consistent with the ventral to dorsal maturation of this structure (Shwartz et al., 2012).

Our analysis did not find any significant morphological abnormalities in the development of major structures of the viscerocranium. These data are consistent with the mild to moderate phenotypes present in humans with *cblC*. Cerone and colleagues reported mild phenotypes that include long face, high forehead, large-low set ears, and a flat philtrum (Cerone et al., 1999). Data derived from the hg13 allele and human patients contrasts with the phenotypes reported in mouse models of *cblC*, which are much more dramatic including facial clefting (Chern et al., 2020). The hyosymplectic which we observed as defective, can be described as a fusion of the hyomandibula with a symplectic rod. It connects to the neurocranium through the anterior region of the otic cartilage and helps to secure the jaw to the rest of the head (Mork and Crump, 2015). The hyomandibula is homologous to the stapes, one of the 3 bones of the inner ear (Mork and Crump, 2015). The palatoquadrate produces homologous structures to the malleus and incus, the other two bones of the inner ear. It remains to be answered whether chondrocyte intercalation defects in the hyosymplectic underlie the low set ears in humans but based on our results, this is a strong possibility.

Abnormal craniofacial development has also been observed in patients with mutations in *HCFC1* and *THAP11*, both of which are upstream regulators of *MMACHC* expression. In fact, knockdown of both genes in the zebrafish and mouse results in craniofacial abnormalities (Chern et al., 2022; Quintana et al., 2017b, 2014). Interestingly, exogenous expression of *MMACHC* can restore facial development after loss of *hcfc1* in zebrafish, but not in the mouse (Chern et al., 2022; Quintana et al., 2014). Restoration in mouse was performed by expressing *MMACHC* specifically in NCCs, whereas rescue in zebrafish was performed through microinjection at the single cell stage, a global expression strategy. This is an interesting difference because we do not find abnormal expression of early NCC markers in the hg13 allele. Moreover, the mouse models of HCFC1 and THAP11 are associated with increased metabolite accumulation, but the early zebrafish development is not (Quintana et al., 2014). Hence the function of MMACHC in the craniofacial development in mice and zebrafish could be quite different. A previous study also noted a maternal effect associated with restoration of facial phenotypes in a *cblC* mouse model (Chern et al., 2020). Zebrafish are likely to have a large maternal effect, as they are externally fertilized.

A function for MMACHC in the transport and enzymatic modification of cobalamin has been clearly established (Froese et al., 2012). But in addition to its enzymatic function, the C-terminal region of the MMACHC protein shows high similarity to TonB domain containing proteins. TonB proteins are outer membrane proteins in bacteria that are known to be involved in energy transduction, cobalamin uptake, and protein interactions with other cobalamin associated proteins (Fofou-Caillierez et al., 2013). As we discussed previously, other inborn errors of cobalamin metabolism have mild craniofacial phenotypes but don’t have mutations in *MMACHC* (Bassim et al., 2009; Coelho et al., 2012, p. 4; Gérard et al., 2015, p. 1; Oladipo et al., 2011). Therefore, we cannot exclude that other cobalamin related proteins interact with MMACHC, perhaps via the TonB domain, and regulate facial development as a unit. Thus, there is much we do not yet understand about the function of MMACHC, particularly beyond its cobalamin binding activity. We noted that the phenotypes in *cblC* can be heterogeneous and the presence of specific phenotypes such as the craniofacial phenotypes do not always correlate with the severity of cobalamin defects (Lerner-Ellis et al., 2009). This is also true in the context of *cblX*, where the severity of metabolic phenotypes is variable (Yu et al., 2013). Because of this, we sought to directly test the function of cobalamin binding. We performed a restoration experiment by injecting mRNA encoding the patient derived p.Gly147Asp variant. The p.Gly147Asp variant is present in the cobalamin binding domain (amino acids 122-156) (Liu et al., 2010) and does not bind hydroxycobalamin or cyanocobalamin (Froese et al., 2009), consequently disrupting cobalamin metabolism. Expression of the p.Gly147Asp encoding mRNA fully restored chondrocyte organization providing evidence that cobalamin binding was not required for proper chondrocyte development. Thus, it is likely that the mild to moderate facial phenotypes present in patients and the hg13 allele are the result of an unknown, but cobalamin independent function.

In summary, we provide functional analysis of *MMACHC*, a gene that is mutated in *cblC* and has been implicated in craniofacial development. We observed a fully penetrant chondrocyte intercalation defect in the hyosymplectic cartilage, although early NCC development was normal. These deficits were restored by patient variant of MMACHC unable to bind cobalamin derivatives. Our data provides new insight into the function of MMACHC and suggests a putative function for the protein which has yet to be fully characterized.

## Acknowledgements

The authors would like to expressly thank Dr. Charles Venditti for providing the *hg13 mmachc* allele. We would also like to thank Rahab (Scarlett) Jaramillo, Valeria Virrueta, Jennifer Davila, and Yahir Davila for technical expertise related to zebrafish husbandry and genotyping.

## Data Statement

Data is available upon request from the corresponding author.

## Abbreviations

cblC: methylmalonic aciduria and homocystinuria, cblC type
cblX: methylmalonic aciduria and homocysinuria, cblX type
DPF: days post fertilization
HPF: hours post fertilization MMA-methylmalonic acid
pME-MCS: plasmid middle element-multiple cloning site
pDestTol2CG2: plasmid destination tol2 vector
p5e-Ubiquitin: plasmid 5 element with ubiquitin promoter
p3ePolyA: plasmid 3’ element with polyadenylation tail
min: minute
pGEM T: plasmid TA vector
PBS: phosphate buffered saline
PBT: PBS with 0.1% Tween 20
tRNA: transfer RNA
RNA: ribonucleic acid
DNA: deoxyribonucleic acid
Fire LUT: Fire look up table
M: Meckel’s cartilage
PQ: palatoquadrate
CH: ceratohyal
HS: hyosymplectic

